# Different features of cholera in malnourished and non-malnourised children: analysis of 10-year surveillance data from a large diarrheal disease hospital in urban Bangladesh

**DOI:** 10.1101/460782

**Authors:** Sharika Nuzhat, Md Iqbal Hossain, Nusrat Jahan Shaly, Rafiqul Islam, Soroar Hossain Khan, A S G Faruque, Pradip Kumar Bardhan, Azharul Islam khan, Mohammod Jobayer Chisti, Tahmeed Ahmed

## Abstract

**Background:** Malnourished children are more prone to infectious diseases including severe diarrhea compared to non-malnourished children. Understanding of the differences in the presentation of severe diarrhea such as cholera in children with varying nutritional status may help in the early identification and management these children. However, data are scarce on differences in the presentation in such children. Thus, we aimed to identify the clinical differentials among children with cholera with or without malnutrition.

**Methods:** Data were extracted from diarrheal disease surveillance system (DDSS) of the Dhaka Hospital of icddr,b for the period, January 2008 to December 2017. Among under-five children, cholera positive (culture confirmed) and malnourished children (weight-for-age, weight-for-length or height-for-age Z score (WAZ, WHZ or HAZ) <-2) were considered as the cases (n=305) and children with cholera but non-malnourished (WAZ, HAZ, and WHZ ≥-2.00 to ≤+2.00) were the controls (n=276).

**Results**: A total of 14,403 under-five children were enrolled in the surveillance system during the study period. After adjusting for potential covariates such as maternal illiteracy and slum dwelling, it was revealed that under-five malnourished children with cholera significantly more often presented to the hospital during evening hours (6 pm to 12 mid-night) (OR=1.64, 95% CI=1.16-2.31, P<0.05), had fathers who were illiterate (OR=1.70, 95% CI=1.11-2.62, P<0.05), presented with history of cough within last 7 days (OR=1.64, 95% CI=1.10-2.43, P<0.05), dehydrating diarrhea (OR=1.70, 95% CI=1.15-2.53, P<0.05), and had longer hospitalization (OR=1.50, 95% CI=1.05-2.14, P<0.05).

**Conclusions:** The study results underscore the importance of understanding of the basic differences in the presentation of severe cholera in malnourished children for prompt identification and the subsequent management of these children. These observations may help policy makers in formulating better case management strategy.

**Author Summary:** Malnourished children are more vulnerable to infectious diseases including cholera in comparison to the non-malnourished children. They often have suboptimal immune function, though there is no precise information on whether there is any difference in associated factor(s) or clinical course of cholera in under-five children with varying nutritional status. Therefore, this study was conducted to elucidate these insights by using the surveillance data of the Dhaka hospital of icddr,b. Among all the under-five children with cholera, 305 malnourished (WAZ or WLZ or HAZ <-2) children constituted as the cases (malnourished), and another 276 non-malnourished (WAZ, HAZ, and WHZ ≥-2.00GtoG≤+2.00) cholera children formed the comparison group.

In this study we revealed that care seeking at evening time was more common in the malnourished children with cholera compared to those without malnutrition. Dehydrating diarrhea was about two folds higher and prolonged hospitalization was frequent in malnourished children with cholera than their counterparts. These key findings may help policy makers in formulating better case management strategy in the near future.

## Introduction

Cholera is a leading public health concern globally, with an estimated 1.3-4.0 million cases occurring each year, worldwide [1]. Moreover, a significant disease burden of cholera has been reported in young children [2]. Higher rates of malnutrition among pre-school children have been observed in Bangladesh [3]. Malnourished children are at a higher risk of severity of diarrhea and death, moreover, the disease severity has been found to be associated with nutritional status, body size, and etiologic agents of diarrheal episodes [4]. A study in Brazil reported that the clinical presentations of early childhood severe diarrhea may vary because of diverse etiologies. Understanding of the differences in the presenting features of severe diarrhea especially in cholera, that are associated with varying nutritional status of the young children is thus critically important for early identification and management these children. Hence, with an attempt to address the existing knowledge gap as well as to share research findings with policy makers for formulating better case management strategy we undertook this study to examine the clinical feature differentials among children infected with *Vibrio cholerae who presented with or without* malnutrition.

## Methods

### Ethical statement

For this study, data were extracted from the electronic database of hospital-based diarrheal disease surveillance system (DDSS) of Dhaka Hospital of icddr,b. The DDSS has the approval from institutional review board of icddr,b (Research Review Committee and Ethical Review Committee) for data analysis. ERC was also pleased with the voluntary participation, maitainance of rights of the participants and confidential handling of personal information by the hospital doctors and accepted this consenting procedure. At the time of enrolment into DDSS, verbal consent was obtained from the parents or the attending caregivers of each child following hospital policy. The verbal consent was recorded by keeping a check mark in the questionnaire that was again assured by showing the mark to parents or caregiver. DDSS is a routine ongoing surveillance in hospitals of icddr,b located in Dhaka, Bangladesh. At the time of consenting, parents or caregivers were assured of ‘any risk being no more than minimal risk’, ‘their participation is voluntary’, ‘their rights to withdraw from the study, and ‘the maintenance of strict confidentiality of disclosed information. They were also informed about the use of collected data for analysis and using the results for improving patient care, conducting researches and also publication without disclosing the name or identity of their children.

### Study population and study site

Dhaka Hospital of icddr,b provides care and free treatment to around 150,000 diarrheal disease patients each year and about 62% of them are children less than 5 years of age. The DDSS systematically (from every 50^th^ patient according to their hospital ID number) collects information including age, sex, socio-demographic characteristics, clinical features, and etiology of diarrhea. Parents or caregivers are interviewed by research assistants who collect demographic, socioeconomic and clinical data. A physician documents the clinical findings including dehydration status. A fresh stool sample is collected and submitted to laboratory for microbiological evaluation. All relevant information is recorded into the electronic database as soon as possible. For the present study, analysis was limited to under-five children who were cholera positive and enrolled into the DDSS from January 2008 to December 2017. Nutritional status of these children was assessed at the time of discharge from the hospital. Weight was measured nearest to 100 g using a digital scale and length/height was estimated using a locally manufactured length board with a precision of 0.1 cm. Nutritional status was assessed by Z-scores following WHO 2006 growth standards.

### Study design

A case control study design was followed. The study group (cases) comprised of malnourished diarrheal children with associated *Vibrio cholerae* infections and those presented at the same time without malnutrition constituted the concurrent comparison group (controls).

### Definition

Malnutrition was defined in children 0-59 months of age with any of the indices of malnutrition such as: weight for age z-score (WAZ) or height for age z-score (HAZ) or weight for height z-score (WAH) < −2. Non-malnourished children had WAZ, HAZ, and WHZ ≥-2.00 to ≤+2.00. A child with cholera had growth of *V. cholerae* in the fecal specimen.

### Data Analysis

Data were analyzed using SPSS for windows (version 20; SPSS Inc, Chicago, IL) and Epi Info 7. Differences in the proportion were compared by the Chi-square test. A probability value of < 0.05 was considered as statistically significant. Strength of association was determined by calculating odds ratios (OR) and their 95% confidence intervals (CI). Logistic regression was performed to identify factors that were considered significantly associated with malnourished cholera children after adjusting for potential confounding variables. Multicollinearity between independent variables was also checked before constructing logistic regression models having a variance inflation factor (VIF) of < 3.0.

## Results

A total of 14, 403 under-five children were enrolled in the DDSS during the study period and 581 children were found to have culture proven cholera. Out of them, according to the eligibility criteria 305 belonged to the study group (cases) while the rest 276 constituted the comparison group (controls). Bi-variate analysis revealed that the cases more often had illiterate mothers and lived in slum settlements compared to the controls. The cases compared to their counterpart commonly reported to the facility at evening hours (6 pm −12 mid-nights), often had history of cough within the last seven days, and were found to seek out care for dehydrating diarrhea. The cases often required longer hospitalization than the controls (Table 1).

**Table 1.** Clinical findings of cholera children with or without malnutrition (2008-2017)

| Variable | Malnourished<br>cholera children<br>(n=305) (%) | Non-malnourished<br>cholera children<br>(n=276) (%) | OR | 95%CI | P value |
| --- | --- | --- | --- | --- | --- |
| Male | 186 (61%) | 169 (61.2%) | 0.99 | 0.71-1.38 | 0.981 |
| Illiterate mother | 105 (34.4%) | 70 (25.4%) | 1.55 | 1.08-2.21 | **0.022** |
| Illiterate father | 117 (38.4%) | 68 (24.6%) | 1.9 | 1.33-2.72 | **0.001** |
| Slum area | 41 (13.4%) | 22 (8%) | 1.8 | 1.04-3.10 | **0.047** |
| Non-sanitary Toilet | 68 (22.3%) | 51 (18.5%) | 1.27 | 0.84-1.90 | 0.300 |
| Use of untreated drinking water | 197 (64.6%) | 161 (58.3%) | 1.3 | 0.93-1.82 | 0.143 |
| No use of antibiotic at home | 115 (37.7%) | 110 (39.9%) | 0.91 | 0.65-1.28 | 0.656 |
| Presence of vomiting | 265 (86.9%) | 237 (85.9%) | 1.1 | 0.68-1.8 | 0.814 |
| Presence of fever | 15 (4.9%) | 9 (3.3%) | 1.53 | 0.7-3.57 | 0.427 |
| Presence of abdominal pain | 87 (28.5%) | 83 (30.1%) | 0.93 | 0.65-1.33 | 0.750 |
| History of cough within last 7 days | 93 (30.5%) | 57 (20.7%) | 1.69 | 0.33-14.29 | **0.009** |
| History of measles | 11 (3.6%) | 12 (4.3%) | 0.82 | 0.36-1.9 | 0.807 |
| Diarrheal duration < 24hours | 129 (42.3%) | 135 (48.9%) | 0.77 | 0.55-1.06 | 0.129 |
| Dehydration (some/severe) | 242 (79.3%) | 184 (66.7%) | 1.9 | 1.32-2.8 | **0.001** |
| Watery stool | 291 (94.4%) | 267 (96.7%) | 0.7 | 0.3-1.63 | 0.543 |
| Use of IV fluid | 145 (47.7%) | 123 (44.7%) | 1.13 | 0.81-1.56 | 0.527 |
| Reporting time<br>(6.00 pm to 12.00 mid-night) | 76 (24.9 %) | 41(14.9 %) | 1.89 | 1.24 -2.88 | **0.004** |
| Length of stay > 24hours | 143(47.4 %) | 96(36 %) | 1.14 | 1.14-2.24 | **0.008** |
| Death | 0 (0%) | 1 (0.4%) | - | - | 0.948 |

% denotes percentage of cholera positive malnourished and non-malnourished group until
mentioned otherwise

Logistic regression analysis adjusting for potential covariates such as maternal illiteracy, and slum dwelling revealed that malnourished children under 5 years of age with cholera significantly more often had paternal illiteracy, history of cough, reported to the hospital in evening hours, presented with dehydrating diarrhea, and had longer stay at hospital (Table 2). Another logistic regression revealed no significant association between untreated drinking water, non-sanitary toilet use, febrile illness and cholera in malnourished children.

**Table 2.** Logistic regression analysis to find out independent factors associated with cholera in malnourished children

| **Characteristic** | **Adjusted OR** | **95%CI** | **P value** |
| --- | --- | --- | --- |
| Illiterate mother | 0.98 | 0.63-1.54 | 0.943 |
| Illiterate father | 1.74 | 1.13-2.67 | **0.012** |
| Slum dwelling | 1.75 | 0.97-3.21 | 0.071 |
| History of cough within last 7 days | 1.61 | 1.08-2.39 | **0.019** |
| Reporting time<br>(6.00 pm to 12.00 mid night) | 1.76 | 1.14-2.73 | **0.011** |
| Dehydration (some/severe) | 1.67 | 1.13-2.48 | **0.011** |
| Length of stay > 24hours | 1.51 | 1.06-2.15 | **0.022** |

## Discussion

The present study observed different features of cholera in children with or without malnutrition. The most important significant observation is the 67% excess risk of dehydrating diarrhea in malnourished cholera children than non-malnourished cholera children. Other important observations in malnourished cholera children with dehydrating diarrhea than non-malnourished cholera children were: i) most often care seeking at evening-night hours, and iii) longer hospitalization of malnourished cholera children.

As expected in malnourished children, lack of immune responses particularly the reduced secretory IgA levels in the gut mucosa along with hypochlorhydria or achlorhydria made these study children more vulnerable to diarrheal illnesses with relatively lower inoculums. Possible explanation for more dehydrating diarrhea in malnourished cholera children are: these children are more often slum dwellers with poor water-sanitation and hygienic practices that might have caused the ingestion of larger inoculums of *Vibrio cholerae* resulting in greater challenging dose of cholera toxin. Moreover, since malnourished children are likely to have an increased area of gut mucosal surface compared to their body weight than the non-malnourished children they are more vulnerable to higher purging rate and resultant greater stool output during diarrhea [4]. In case of malnourished children with cholera, slower turnover rate of gut mucosal cells, deficiencies of intestinal enzymes, micronutrients, and impaired immune responses with exposures to larger inoculums because of their dwelling in more contaminated environments in the slums might have caused more severe disease and delayed recovery, thereby, longer hospitalization.

Dewan *et al.* reported that children with associated *Vibrio cholerae* infections were significantly more severely underweight, stunted, and wasted. The study indicated that such association may be due to impaired gastric barrier, hypochlorhydria, and prolonged intestinal mucosal injury that are commonly observed in malnourished children [5]. A study in urban Bangladesh evaluated the role of common diarrheal pathogens and revealed that children with *Vibrio cholerae* infections were 5.5 times more likely to be associated with dehydrating diarrhea than that in case of children without this causative agent [6] and these children were significantly more malnourished. Similar to our findings, other cholera researchers from Bangladesh revealed marked prolongation of duration of diarrhea in undernourished children [7]. Exerting greater emphasis on stool output, Palmer *et al.* indicated that higher inoculum size is likely to cause greater intestinal mucosal surface involvement causing higher stool volume per unit of time than a longer duration of diarrhea [7]. On the basis of a study in the rural settings of Bangladesh, Black *et al.* reported that child’s small body size because of young age and low nutritional status are more likely to result in more fluid loss (per kg body weight) during diarrhea and such children are more vulnerable to severe dehydration and death if not properly treated by appropriate rehydration therapy [4]. Most studies that reported association between malnutrition and severity of diarrhea did not take into consideration the role of enteropathogens in causing severity of dehydration and longer duration of the episode [5, 8]. Another study in Brazil reported that children with fever, vomiting or both would capture 75% of the children at risk for dehydrating diarrhea [9]. However, this study compared the inpatient cases versus outpatient controls without relating to the etiology and nutritional status. In our study particularly with cholera positive children, vomiting (86.9% vs 85.9%), fever (4.9% vs 3.3%) or abdominal pain (28.5% vs 30.1%) did not demonstrate any significant difference in cholera children with or without malnutrition.

Victora *et al*. observed strong association between low body weight of infants (regardless of age) and risk of higher dehydration. Low body weight was observed to be a superior determinant in comparison to the anthropometric indices for predicting dehydrating diarrhea in children reporting to the health facility. The study mentioned that children with low body weight are young, malnourished or both. These children have larger gut surface compared to their body size in addition to greater purging rate due to diarrhea as compared to older children [8]. Thus malnourished cholera children during hospitalization require intensive treatment with adjunct appropriate antimicrobial and zinc therapy, careful assessments of dehydration at intervals, appropriate dietary intervention with closer follow-up. Malnourished children are more prevalent in families with low income and poor housing along with compromised water-sanitation and hygienic practices. These children should be targeted for health education at household level along with support for continued breast feeding, initiation of rehydration therapy soon after the onset of diarrheal illnesses to prevent severity of disease with early referral of dehydrating children to appropriate facilities to avoid unnecessary death.

These malnourished cholera children were mostly from the urban slums and due to the nature and working hours of the father or mother or both, they presented to the facility during evening hours or later. Another important observation from our study was that the malnourished cholera children had a greater frequency of a history of cough within the last seven days. Cough is one of the key clinical features of respiratory tract infections. The most common infections for malnourished children are gastrointestinal and respiratory infections [10]. The first line of defense mechanism for these infections is the innate immunity, particularly the epithelial barriers and the mucosal immune response [11]. Malnourished children significantly suffer from compromised mucosal barriers of the gastrointestinal, respiratory and urogenital tracts.

The study may be replicated in other geographical and cultural settings to see if the same clinical features play a similar role in causing dehydrating diarrhea in malnourished cholera children. Cholera outbreaks in emergency settings, such as settlements for displaced population and in the aftermath of natural calamities, are now commonly encountered. In such situations, the treatment of children with severe acute malnutrition and cholera is difficult in terms of both competences of clinicians well as coordination of logistics. The findings of our study are likely to be very helpful in such situations emphasizing the critical need in keeping SAM children with cholera for a longer period of time in the diarrhea treatment center before their referral to a nutritional rehabilitation or outpatient treatment center.

This study was conducted in an urban hospital and vast majority of the patients represented with poor socio-economic background. Our study children had higher degree of infection that required hospitalization because of severe illness and they represented a relatively small proportion of children while the vast majority of children with less severe disease received care at the household level and did not seek care from the present facility. Respondents were mothers who presented to the facility with their child; 27% had no formal schooling and 60% of their children were malnourished and those who presented from urban slums, 65% were malnourished. Thus our study children might not be representing the greater population. However, our results are likely to generate several hypotheses. Future studies could better describe the changes in presenting features along with etiology specific changes in clinical features in malnourished children over time period. Along with the unbiased systematic collection of data, a larger sample size, high quality laboratory performance and use of probing techniques in interviewing of mothers or caregivers had thus been the strengths of the study.

## Acknowledgments

This research study was funded by core donors which provide unrestricted support to icddr,b for its operations and research. Current donors providing unrestricted support include: Government of the People’s Republic of Bangladesh; Global Affairs Canada (GAC); Swedish International Development Cooperation Agency (Sida) and the Department for International Development (UK Aid). We gratefully acknowledge these donors for their support and commitment to icddr,b's research efforts.

